# reanalyzerGSE: tackling the everlasting lack of reproducibility and reanalyses in transcriptomics

**DOI:** 10.1101/2023.07.12.548663

**Authors:** José L Ruiz, Laura C Terrón-Camero, Julia Castillo-González, Iván Fernández-Rengel, Mario Delgado, Elena Gonzalez-Rey, Eduardo Andrés-León

**Affiliations:** Institute of Parasitology and Biomedicine “López-Neyra”, CSIC (IPBLN-CSIC), 18016 Granada, Spain

## Abstract

**Summary:** In the current context of transcriptomics democratization, there is an unprecedented surge in the number of studies and datasets. However, advances are hampered by aspects such as the reproducibility crisis, and lack of standardization, in particular with scarce reanalyses of secondary data. reanalyzerGSE, is a user-friendly pipeline that aims to be an all-in-one automatic solution for locally available transcriptomic data and those found in public repositories, thereby encouraging data reuse. With its modular and expandable design, reanalyzerGSE combines cutting-edge software to effectively address simple and complex transcriptomic studies ensuring standardization, up to date reference genome, reproducibility, and flexibility for researchers.

**Availability and implementation:** The reanalyzerGSE open-source code and test data are freely available at both https://github.com/BioinfoIPBLN/reanalyzerGSE and 10.5281/zenodo.XXXX under the GPL3 license.

**Supplementary data** are available.

## INTRODUCTION

Latest technological advances now allow laboratories to sequence any sample with relatively low investment. Transcriptomics is one of the most popular -omics and researchers routinely apply RNA-seq-related techniques to estimate gene and transcript expression [1]. However, while wet-lab protocols have been greatly democratized and the field jumps at the so-called single-cell and spatial revolutions [2-5], average users generally face major challenges due to the lack of computational expertise to perform data analyses. Bioinformatics is up to the challenge and many statistical approaches and protocols have been popularized [6, 7]. However, despite remarkable efforts, their performance may be variable and all are far from being a standard all-in-one solution, with methods being adjusted almost on a case-by-case basis [8-12]. In the current context of reproducibility crisis [13-18], published results are also typically incorporated in following studies, but new hypotheses would often require updating reference genome and annotation and fine-tuned reanalyses, which are rare. Therefore, while deposited in public databases that increasingly try to facilitate the accessibility and use of raw data [19-21], countless transcriptomics datasets are largely unexplored beyond their original publication (*i*.*e*., remain restricted). This is crucial because the reuse of data is one of the most important FAIR principles that should sustain the new era of Open Science [22], particularly regarding the field of Bioinformatics [23, 24]. Overall, there is an urgent need for standardization to address several issues hampering transcriptomics that can lead to errors in biological interpretations and conclusions. These may include: 1) the use of obsolete genomic references [25, 26], 2) the presence of contamination (*e*.*g*. microorganisms [27] or rRNA [28]), 3) the use of inappropriate software or statistical approaches (*e*.*g*. overlooked covariables and confounders [29], unexpected batch effects [30], variable filtering of noise or low expression [31-33], differences in quantification [8], normalization [34], pre-processing [35], differential gene expression [31, 36, 37], or functional enrichment [38-40]), and 4) diverse errors (*e*.*g*. insufficient reporting [41], errors in gene naming errors [42, 43], complex software [44-46], or data handling [47])

To overcome these, amongst others, we introduce here reanalyzerGSE, an easy-to-use pipeline capable of addressing full RNA-seq and microarrays analyses in an automatic and standardized manner. From raw transcriptomics data and with minimum user input, ready-to-visualize reports are generated to be directly interpreted by any non-expert user. The processing steps integrate dozens of tools in both mandatory and optional steps to achieve: 1) reorganization of local data or comprehensive download of raw reads and metadata from multiple public repositories following naming conventions, 2) subsampling of data and normalization, 3) quality control, contamination and adapter removal, 4) alignment to up-to-date reference sequences, 5) quantification and correction of batch effects, 6) flexible filtering of noise, 7) differential gene expression analyses, 8) functional enrichment, clustering and network analyses, and 9) generation of interactive human-readable results.

reanalyzerGSE has been tested with datasets of organisms spanning different domains of life and technologies, including complex transcriptomic studies requiring unusual processing. As a case example, we used cortistatin, which is a neuropeptide with a prominent immunomodulatory role [48]. Despite its biological relevance, it is typically discarded as noise by the vast majority of software, due to its extremely low level of expression. To sum up, our aim is to facilitate transcriptomics data computation to both novel and expert users, enabling straight-forward implementation, reproducible and standardized analyses of primary data, and routine reanalyses of secondary data. Our pipeline is modular, parallelizable and scalable, and outperforms alternatives.

## MATERIAL AND METHODS

### Implementation details

reanalyzerGSE is implemented as a Bash pipeline integrated with multiple external software, including R and Perl scripts. The universality of Bash makes possible that researchers from any background can use or adapt tools easily, without the more specific knowledge required by more recent workflow management options. However, we implemented abundant Bash-based solutions and workflow rules to ensure modularity and compatibility systems-wide, such as 1) basing all execution on one-liners, 2) automatically setting default values to non-experienced users, 3) automatically resuming any interrupted run from the appropriate step, 4) when possible automatically performing RAM management and parallel processing (via GNU parallel [49]) and RAM management to make the most of any resources available, 5) creating a fine-tuned structure of folders, results and log files to guarantee the traceability of data, errors and resources, 6) dealing with all steps and saving all intermediate results independently to allow for compatibility with any other software or downstream analyses, or 7) achieving installation of dependencies effortlessly by a one-liner wrapper leveraging the conda package manager with isolated distributable environments and frozen versions that avoid dependencies conflicts, conforming to latest recommendations in the field [46, 50, 51].

Our pipeline has been designed to be versatile and customizable, offering at the same time standardization, reproducibility and flexibility. reanalyzerGSE supports all the formats available for raw sequencing data, both locally available or public in a plethora of databases (*e*.*g*. GEO, SRA, ENA, DDBJ…). This would allow users to routinely reanalyze available secondary datasets, instead of directly integrating results (*i*.*e*. expression levels) by others, which may be inaccurate or erroneous in the context of different or tailored studies. While not recommended, users may also skip pre-processing and alignments to only perform downstream analyses on already-available counting or expression matrices. An overview of the full and reliable transcriptomic analyses automatically performed by our pipeline is: 1) Data loading (*i*.*e*., gathering local data or downloading from public databases), 2) data pre-processing (*i*.*e*. decontamination, rRNA removal or downsampling), 3) data processing with alternatives, 3.1) full microarray pipeline, 3.2) summary of average profiles in single-cell data, 3.3) full RNA-seq pipeline (*i*.*e*. strandedness prediction, quality control, alignment, quantification, variable noise filtering, normalization, batch effect correction, replicability analyses), 4) differential gene expression analyses, 5) functional enrichment, network or clustering analyses, 6) plotting of results and transcriptomic profiles of genes of interest. The pipeline also detected whether the input data are any kind of RNA-seq or microarrays. Given the extensive requirements of a full scRNA-seq approach, if a reanalysis of a scRNA-seq dataset in a public database is requested, reanalyzerGSE will automatically look for the already-available counts and offer a restricted analyses of the collapsed counts in a bulk-like manner. Figure 1 shows the summarized flowchart of our pipeline, and a more detailed description of the features and advantages of reanalyzerGSE when compared to other alternatives is available in Supplementary Table 1.

**Figure 1:**
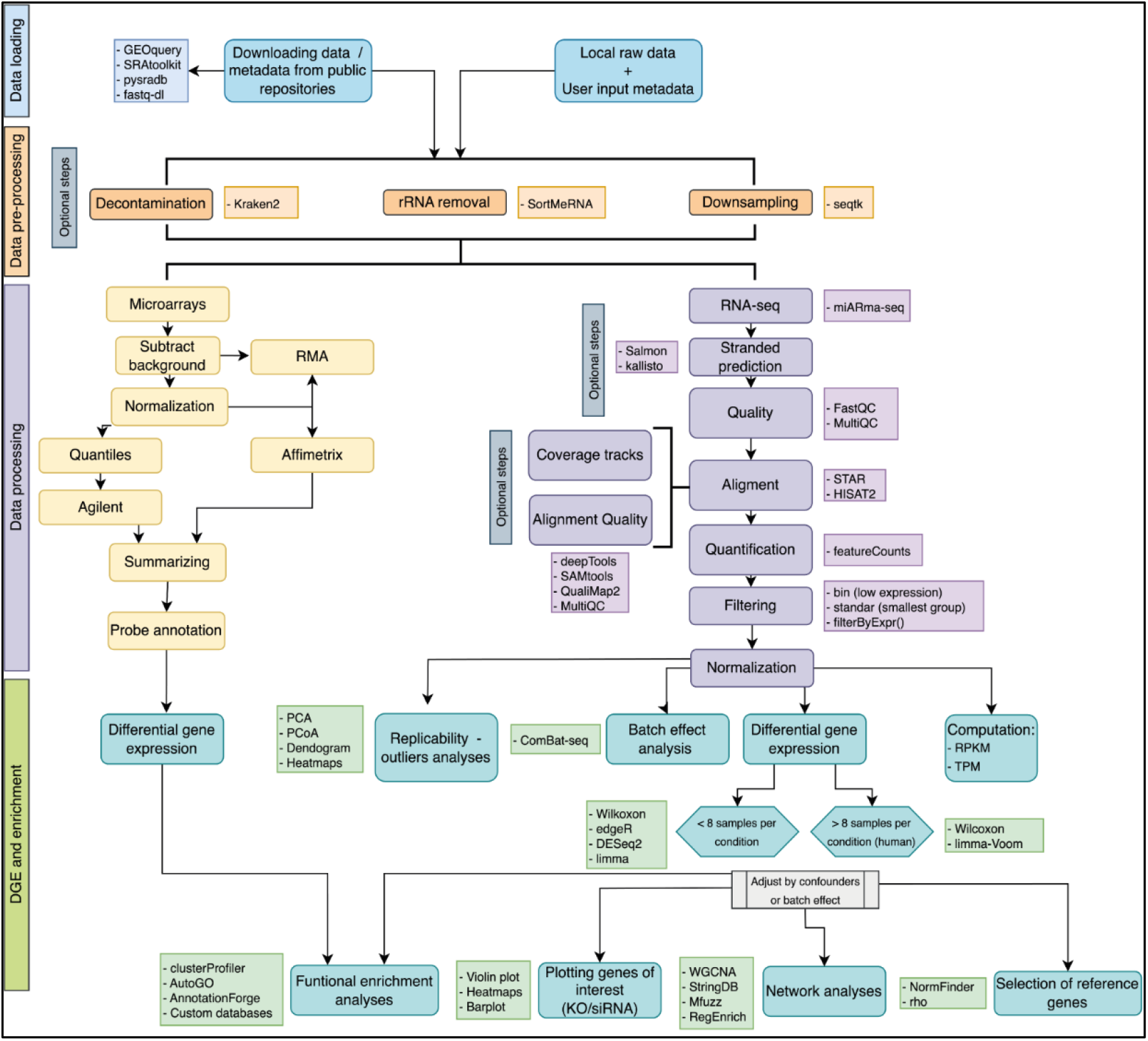
reanalyzerGSE workflow with the most relevant steps and software highlighted at each step.

Notably, we have implemented up-to-date solutions anywhere possible to ensure proper processing, such as a novel statistical approach for larger large-sample differential gene expression analyses [37], a recent framework to avoid confounders in functional enrichment analyses [38, 52], or an extensive update of the miARma-seq suite for RNA-seq analyses [53, 54].

## RESULTS

The primary aim of reanalyzerGSE was to make possible that non-expert users perform reliable and reproducible analyses of transcriptomics data. To that end, we followed the most popular recommendations to create a full RNA-seq/microarrays workflow [35, 46, 51]. When compared to others, Supplementary Table 1 shows that our tool is one of the most helpful and complete available, providing ready-to-interpretate biological results via integrated functional analyses and exploratory analyses and graphics any genes of interest provided by the user. Thanks to its iterative implementation, reanalyzerGSE is also fast, with a performance that allows a user to run it iteratively, fine-tuning any parameter required after preliminary exploratory analyses. For instance, our pipeline performed the full analysis of a mouse dataset comprising 6 samples of oligodendrocytes (total of 7.7G bases, GSE118451) [55] in 40 minutes, and it scaled well, as it fully reanalyzed a larger study of 68 samples of endothelial cells from disease models (total of 750.2G bases, GSE95401) [56] in 21 hours (Supplementary Figure 1 and Supplementary Table 2).

**Supplementary Figure 1:**
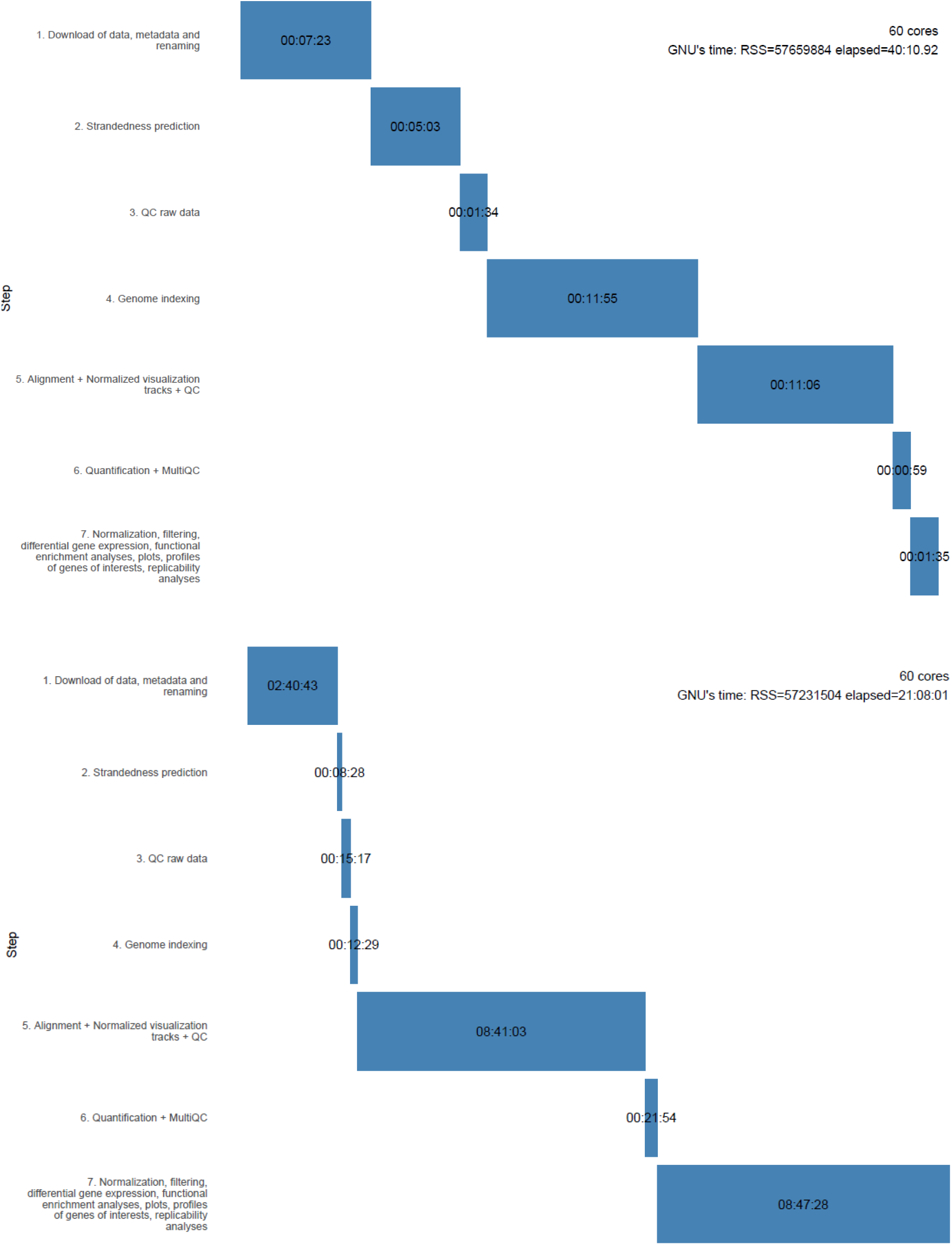
Gantt charts depicting the runtimes for each one of the main steps of the reanalysis of raw data by two studies/entries for GEO: GSE118451 (top) and GSE95401 (bottom). The time require by each step is shown in the format hours:minutes:seconds. The output of GNU’s time after the processing with 60 cores is shown (RSS and elapsed show the total RAM and processing time, respectively).

Beyond securing robustness and completeness of typical use cases of transcriptomics, when developed reanalyzerGSE we focused on those processing steps that may be typically associated with non-trivial errors that lead to inconsistent results or biological interpretations, particularly to non-expert users. For example, filtering out the genes with low counts beforehand is an important step to improve statistical power in differential expression analyses, because low count genes will have more statistical noise and more variation in expression, being less likely to pass the significance threshold in the differential expression and producing large and inaccurate Fold Change values [31, 33]. Therefore, they would be contributing to the number of multiple tests and affecting calculation of false discovery rate (FDR), while not contributing to the number of truly expressed or differentially expressed genes (DEGs). To address this point, users typically discard genes below an arbitrary threshold of raw counts, which is typically not reported, or use obscure approaches, such as “independent filtering” by DESeq2 [57] that automatically settles on a threshold that may be variable between studies. To account for this, we added alternative approaches to reanalyzerGSE, such as the filterByExpr function by edgeR [58], a “standard” filter that retains genes with more than 3 c.p.m counts, or a “bin” filter that retains genes if they were assigned more than one count in any sample. In general, genes with biological relevance may display low expression, close to the detection threshold of the techniques. Low expressed genes that are incorrectly discarded as noise may include diverse functions, such as hormones, neuropeptides, tumour microenvironments [59] or responses against radiation [32]. It has also been reported that hundreds of relevant genes functionally-related to human disease are discarded in RNA-seq analyses to measure group-level expression [8]. Here, we show the case example of the neuropeptide cortistatin in a human dataset of stem cell derived astrocytes and brain endothelial cells (https://www.ncbi.nlm.nih.gov/geo/query/acc.cgi?acc=GSE181332). Its expression would be deemed to be negligible if a standard filtering was performed, while our bin filtering would recover its meaningful expression with two of the most popular alignment software (Supplementary Figure 2).

**Supplementary Figure 2:**
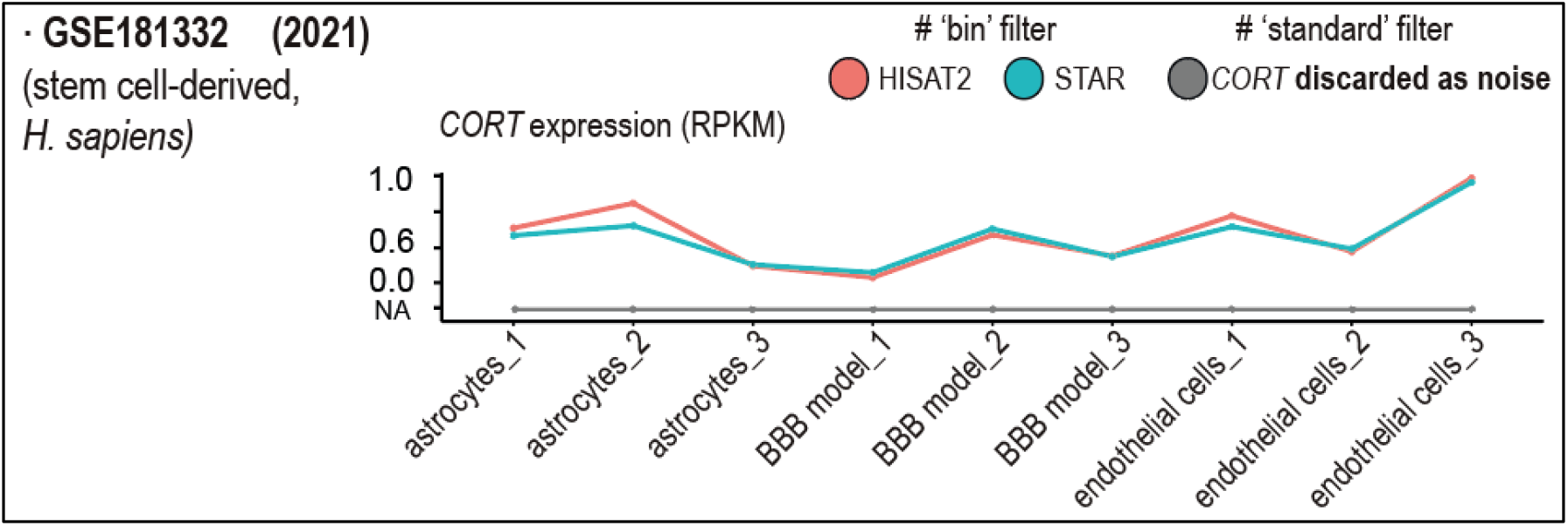
Two reanalyses of the human dataset GSE181332. Lineplots show the expression level (RPKM) of the neuropeptide cortistatin (CORT) is shown with the “standard” filter (in gray, not detected), and the “bin” filter and the aligners HISAT2 and STAR (red and blue color, respectively).

Apart from the tailored filtering, another strength of reanalyzerGSE is rapidly indexing any reference genome provided, archiving it for following runs. This way, our pipeline can be used to easily reanalyse secondary data with alternative versions of the reference sequences, which may be crucial. Supplementary Figure 3 shows the case example of a tobacco dataset used in a study addressing cold tolerance of different cultivar [60]. It can be observed how the number of DEGs, and ultimately the inference of function and biological interpretations, would greatly change in the case of non-model organisms or samples lacking gold-standard genomes, depending on the reference sequences used.

**Supplementary Figure 3:**
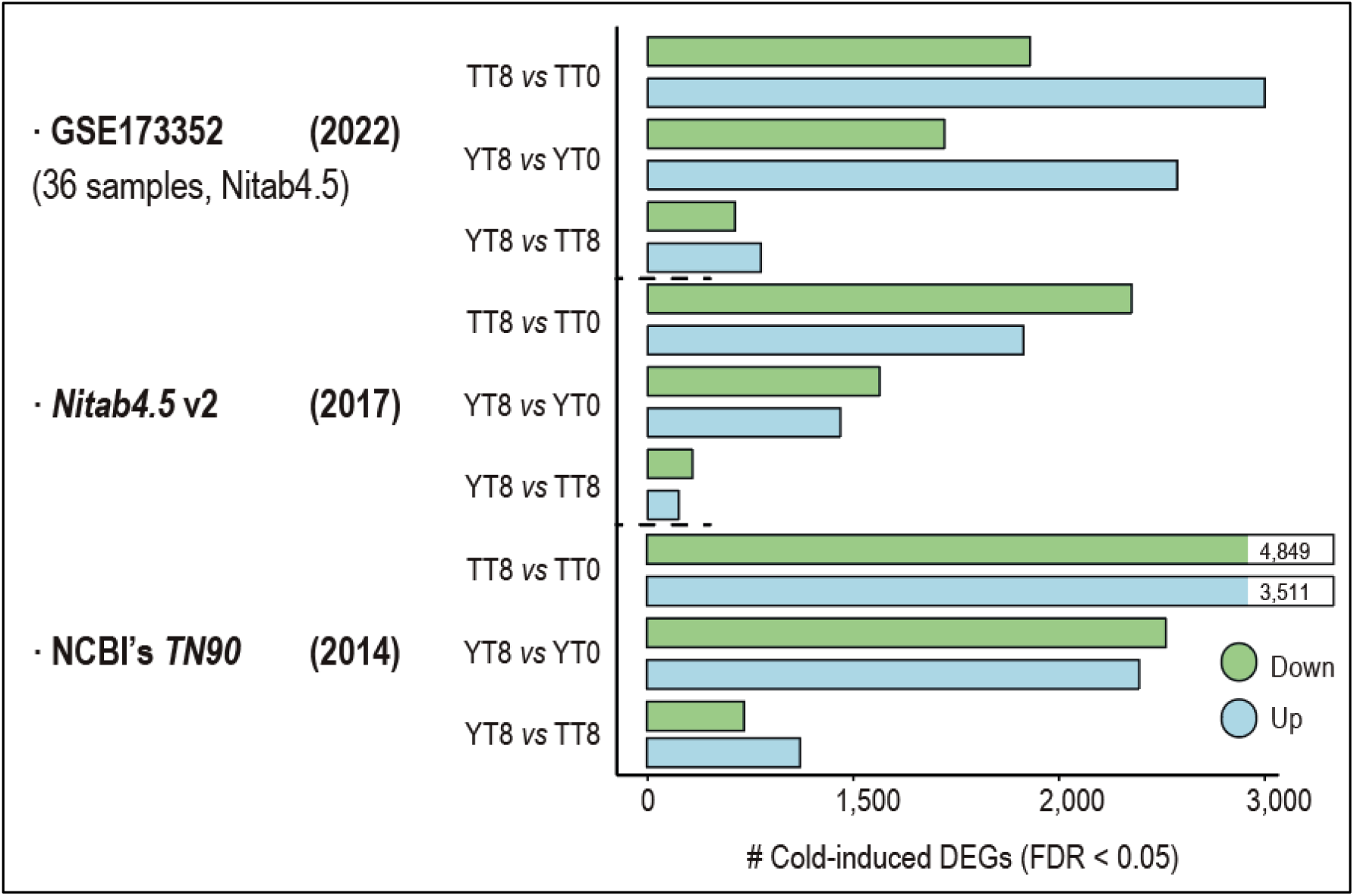
Lineplots show reanalyses of the dataset GSE173352 with three alternative tobacco genomes, including the one used in the original publication, the most recent collapsed reference in chromosomes, and an outdated version that is still the hallmark by NCBI. The number of DEGs is shown for a few comparisons between the two tobacco cultivars (taiyan, TT, or yanyan, YY) and two time points (0 and 8 hours).

## CONCLUSIONS

We present reanalyzerGSE, an automatic pipeline for the analysis of transcriptomics studies (*i*.*e*., RNA-seq, microarrays). It has been implemented using Bash and the conda package manager, offering the largest compilation to date of tools for RNA-seq data analyses. reanalyzerGSE will be actively maintained and in continuous development, incorporating new tools suggested by users and conforming to community standards. Future work will also include extending the functionality to state-of-the art single-cell approaches. The open-source code, together with full documentation and tutorials are available at https://github.com/BioinfoIPBLN/reanalyzerGSE.

## DATA AVAILABILITY

All data are incorporated into the article and its online supplementary material.

